# Analysis and modeling of cancer drug responses using cell cycle phase-specific rate effects

**DOI:** 10.1101/2020.07.24.219907

**Authors:** Sean M. Gross, Farnaz Mohammadi, Crystal Sanchez-Aguila, Paulina J. Zhan, Tiera A. Liby, Mark A. Dane, Aaron S. Meyer, Laura M. Heiser

## Abstract

Identifying effective therapeutic strategies that can prevent tumor cell proliferation is a major challenge to improving outcomes for patients with breast cancer. Here we sought to deepen our understanding of how clinically relevant anti-cancer agents modulate cell cycle progression. We genetically engineered breast cancer cell lines to express a cell cycle reporter and then tracked drug-induced changes in cell number and cell cycle phase, which revealed drug-specific cell cycle effects that varied across time. This suggested that a computational model that could account for cell cycle phase durations would provide a framework to explore drug-induced changes in cell cycle changes. Toward that goal, we developed a linear chain trick (LCT) computational model, in which the cell cycle was partitioned into subphases that faithfully captured drug-induced dynamic responses. The model inferred drug effects and localized them to specific cell cycle phases, which we confirmed experimentally. We then used our LCT model to predict the effect of unseen drug combinations that target cells in different cell cycle phases. Experimental testing confirmed several model predictions and identified combination treatment strategies that may improve therapeutic response in breast cancer patients. Overall, this integrated experimental and modeling approach opens new avenues for assessing drug responses, predicting effective drug combinations, and identifying optimal drug sequencing strategies.

## INTRODUCTION

Developing transformative anti-cancer therapies requires drug combinations^1^, yet rationally identifying effective combination therapy regimens remains challenging^2–5^. Many anti-cancer agents are designed to impact cell proliferation and viability, which suggests that incorporating information about how individual drugs impact cell cycle behavior can lead to improved predictions about drug combination effects. The mammalian cell cycle can be separated into four linked phases (G_1_, S, G_2_, and M) with multiple checkpoints (restriction point, DNA damage checkpoint, and the spindle assembly checkpoint)^6–9^. Cell cycle phases and checkpoints are primarily composed of different molecular entities and, consequently, each phase is regulated in different ways, which results in a minimal correlation between cell cycle phase durations in individual cells^10^. This independence between phases and checkpoints has implications for cancer treatment because many cancer drugs directly target different aspects of the cell cycle; for example, CDK4/6 inhibitors block progression out of G_1_ phase^11^, while the nucleoside analog gemcitabine activates the DNA damage checkpoint by targeting DNA synthesis during S-phase^12^. Together, these findings imply that drug-induced changes to cell numbers can be achieved through distinct cell cycle-dependent molecular mechanisms. For example, these observations suggest that combing two drugs that each reduce the rate of G_1_ progression will lead to deeper reductions in the rate of G_1_ progression, rather than an increase in cell death. Further, this framework predicts dose-dependent impacts: at sub-saturating doses, these G_1_ effects will add together to reduce cell numbers, while at higher saturating doses the cell number will peak at the maximum cytostatic effect. This general idea of drug combination efficacy was recently explored in a study of the multi-drug CHOP protocol used in the treatment of non-Hodgkin Lymphoma, which showed that the effectiveness of this drug combination could be attributed to the fact that each agent had non-overlapping cytotoxic effects^13^. The CHOP protocol also demonstrates the benefit of drug combinations to improve patient outcomes. Considering both cell cycle and cell death effects in greater detail, therefore, has the potential to significantly improve drug combination predictions.

The classic approach to quantifying drug response is to calculate the number of cells 72 hours after drug treatment and assume cells are undergoing exponential growth^14–17^. Other approaches to quantify drug response include compartmental models such as pharmacokinetic and pharmacodynamic (PK-PD) models that consider drug uptake and population dynamics^18^. Recent advances in methodological and quantitative approaches enable assessment of the impact of therapies on cell growth rates, rather than static cell counts^19^, which yields more robust correlations between molecular features and drug sensitivity^19,20^. However, while growth rate approaches significantly improve quantification, they provide limited information about cell cycle effects. A related approach, fractional proliferation, which models the number of cycling, quiescent, and dying cells in a drug-treated population, incorporates growth rates and assumes that cells irreversibly exit the cell cycle into quiescence^21^. Recent studies demonstrate that cells may not irreversibly exit the cell cycle and instead may extend the duration of a specific cell cycle phase before restarting progression through the cell cycle^22^. These prior studies motivate our interest to deeply assess the influence of drugs on specific cell cycle phases and progression through the cell cycle.

In this report, we quantify and incorporate cell cycle phase effects in an analysis of drug responses to single agents and their combinations. We used live-cell imaging of a panel of molecularly diverse breast cancer cells engineered to express a cell cycle reporter and tracked the dynamics of cell number and cell cycle phase in response to single drugs and drug combinations. Across single drugs, we observed distinct cell cycle effects, which led to similar final cell numbers, with phase-specific responses that were oscillatory over time due to the temporal impacts on the cell cycle. To describe these responses, we developed a computational model that uses a linear chain trick (LCT) to account for the delay from cell cycle phase transit time upon drug treatment. This LCT model correctly inferred single drug responses across time as well as the drug-induced oscillatory cell cycle dynamics. We used this model to predict the effect of unseen combinations of drugs that impact different aspects of the cell cycle.

Experimentally testing several drug combinations, we validated that responses were primarily determined by the specific cell cycle effects of each drug pair. These studies reveal the complexity of cell behavior underlying drug responses, provide mechanistic insights into how individual drugs modulate cell numbers and yield a framework for how the combination of different drugs can be rationally modeled and predicted.

## RESULTS

### Drug treatments induce distinct changes in cell number and cell cycle phasing

To track drug responses in individual cells, we genetically engineered HER2+ AU565 breast cancer cells to stably express the HDHB cell cycle reporter^22^ and a nuclear-localized red fluorescent protein (**Fig. 1A,B**). We then treated cells with escalating doses of five drugs commonly used to treat breast cancer, each targeting different cell cycle phases or apoptotic mechanisms (**Fig. 1C**). Cells were imaged every 30 minutes for 96H, and the number of cells in each cell cycle phase and total cell numbers were quantified.

**Figure 1.**
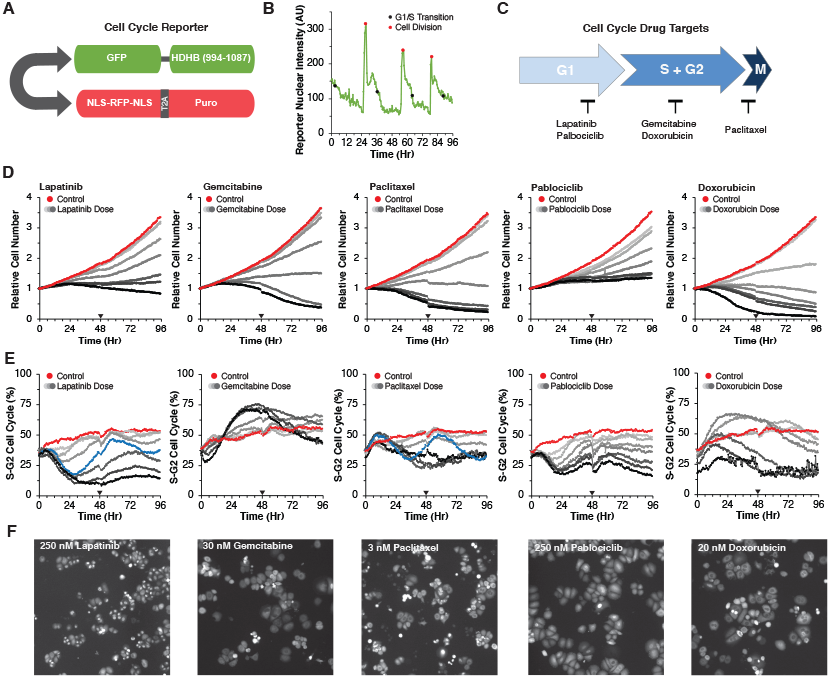
Drugs induce dose- and time-dependent changes in cell cycle behavior. **A**. Schematic of reporter with a bidirectional promoter driving expression of human DNA Helicase B (HDHB) fused to the green fluorescent protein clover, and a second transcript coding for NLS-RFP-NLS, a ribosome skipping domain (T2A), and a puromycin resistance protein. **B**. Quantification of nuclear intensity of the cell cycle reporter in a cell and its progeny across time. The time of G_1_/S transition and cell division are demarcated with black and red circles respectively. **C**. Schematic of the five drugs tested and the cell cycle phase they target. **D**. Average growth curves of AU565 cells tracked every 30 min for 96H across an 8-point dose response for lapatinib, gemcitabine, paclitaxel, palbociclib, and doxorubicin. The null dose is colored red. Line traces show the average from three independent experiments. The black triangle indicates the addition of fresh drug and media. **E**. Percentage of cells in S-G_2_ phase of the cell cycle across doses. 50 nM Lapatinib and 3 nM paclitaxel are colored blue. **F**. GFP images at 39.5H for 250 nM lapatinib, 30 nM gemcitabine, 3 nM paclitaxel, 250 nM palbociclib, 20 nM doxorubicin.

We found that each drug effectively reduced cell numbers in a dose-dependent manner (**Fig. 1D, Sup. Fig. 1**). As expected, paclitaxel, gemcitabine, and doxorubicin led to cytotoxic effects indicated by the final cell numbers dropping below the starting cell numbers (**Fig. 1D**)^23,24^. In contrast, at the highest doses of palbociclib and lapatinib, the final cell numbers were approximately equal to the starting cell numbers, suggesting cytostatic effects. For each drug, the pattern of cell counts varied across time; at high doses responses tended to reach a peak and then decline as the duration of drug exposure increased—an effect most marked for 30 nM gemcitabine where the relative cell number declined from 1.1 at 48H to 0.5 at 96H (**Fig. 1D**)^20,25^.

Next, we sought to identify whether changes in cell numbers arose through the modulation of cell cycle phasing. We observed drug- and dose-dependent changes in the fraction of S-G_2_ cells, which varied over time (**Fig. 1E,F**). For example, lapatinib and palbociclib initially reduced the fraction of cells in S-G_2_ phase in a dose-dependent manner, whereas gemcitabine and doxorubicin initially increased this fraction. Of note, intermediate doses of lapatinib (50 nM) and paclitaxel (3 nM) induced oscillating cell cycle responses, with an initial S-G_2_ reduction near 30H, followed by a second S-G_2_ reduction at 84H. In sum, our approach revealed drug-specific cell cycle changes across time, which confirms that these drugs yield similar final numbers through distinct impacts on the cell cycle.

### A dynamical model of the cell cycle captures the dynamics of drug response

A common approach to model drug effects is to assume exponential growth that varies as a function of drug dose. This approach, although informative, cannot explain the cell cycle dynamics that we observed^26^ and motivated us to develop a dynamical model to capture the observed behavior. As an initial model, we defined a system of ordinary differential equations (ODEs) with transitions between G_1_ and S-G_2_. The parameters of the ODE model were the cell cycle phase progression and death rates, which were assumed to follow a Hill function with respect to drug concentration (**Sup. Table 1**). This model failed to fit the experimental data of G_1_ and S-G_2_ cell numbers (**Sup. Fig. 2**); furthermore, dynamical systems theory dictates that this model is unable to oscillate under any reasonable parameterization^27^.

To address these limitations and capture the observed oscillatory temporal dynamics, we modified our model ‘s assumptions for cell cycle phase durations. We noted that the durations were well described by a gamma distribution and applied the observation that cell cycle phase durations are uncorrelated^10^ (**Sup. Fig. 3A**). Gamma and related distributions model each cell cycle phase as a series of steps, with the key feature that they can model processes wherein there is always some measurable duration before a system (e.g., a cell progressing through the cell cycle) can move to the next state. By fitting the single cell measurements of G_1_ and S-G_2_ phase durations from the untreated control, we estimated the shape parameter of the gamma distribution which determines the number of steps in each phase^28^. This resulted in partitioning the G_1_ phase into 8 and S-G_2_ phase into 20 steps (**Fig. 2A**). We incorporated a “linear chain trick “ into our model, which creates similarly-distributed time delays in the cell cycle phase durations through a mean-field system of ODEs^29^. Additionally, we simplified the model by sharing parameters that were not drug specific, such as the number of cell cycle subphases and the initial fraction of cells in G_1_ phase. We fit all five drug dose responses, varying the drug-specific and shared parameters, simultaneously. Incorporation of this component enabled the model to capture the experimentally observed oscillatory cell cycle behavior and cell cycle phase-specific drug effects. We computed the fitting error of the two modeling frameworks by calculating the sum of squared error of the difference between the data and model predictions across all concentrations and observed that the LCT model had lower error terms (**Fig. 2B**). The fits to lapatinib and palbociclib were particularly improved by the model refinement. Examples of dose-response curves and model fits for lapatinib and gemcitabine are shown in **Figure 2C-H**. Importantly, the model captured the dose-dependent changes to G1 and S-G2 populations as well as the oscillatory dynamics. Estimating the cell cycle phase progression and death rates also enabled calculation of the accumulated amount of cell death across time using inferred cell counts at each phase (**Fig. 2E,H**). The LCT model also performed well for each of the other drugs (**Supp. Fig. 4A-F**).

**Figure 2.**
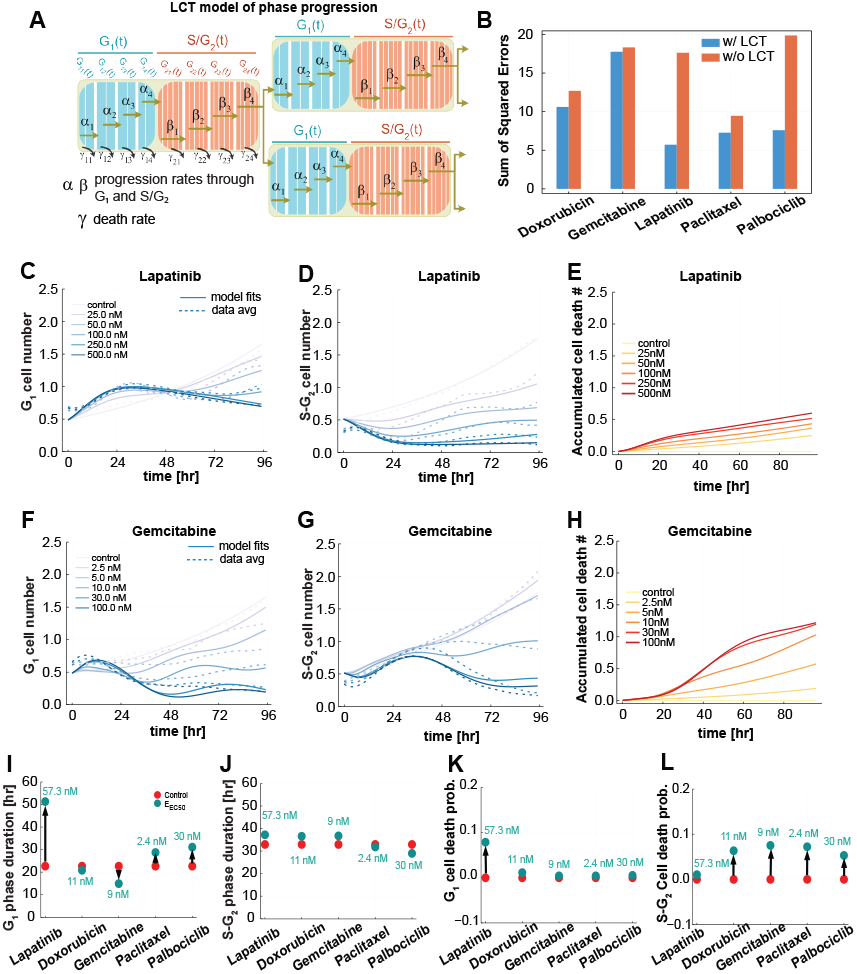
A computational model of the cell cycle captures the dynamics of drug response. **A**. Diagram of the phase transitions in the linear chain trick (LCT) model. *α* _1_, *α* _2_, *α*_3_, and *α* _4_ are the progression rates through G_1_ phase; *β*_1_, *β*_2_, *β*_3_, and *β*_4_ are the progression rates through S-G_2_ phase. Similarly, *γ*_11_, *γ*_12_, *γ*_13_, and *γ*_14_ are the death rates within the G_1_ phase parts, and *γ*_21_, *γ*_22_, *γ*_23_, and *γ*_24_ are death rates within S-G_2_ phase parts. **B**. The sum of squared errors for the fits of each of five drugs over all concentrations with and without the LCT modification. **C-D**. G_1_ and S-G_2_ cell numbers over time, respectively, for lapatinib-treated cells at 5 concentrations and untreated control (solid lines), overlayed with the average of three experimental replicates (dashed lines). **E**. The predicted accumulated dead cells over time for untreated and lapatinib-treated cells at 5 concentrations. **F-G**. G_1_ and S-G_2_ cell numbers over time, respectively, for gemcitabine-treated cells at 5 concentrations and untreated control (solid lines), overlayed with the average of three experimental replicates (dashed lines). **H**. The predicted accumulated dead cells over time for untreated and gemcitabine-treated cells at 5 concentrations. **I-J**. The average phase durations in G1 and S-G2 phases for all five drug treatments. The arrow shows the shift from the control condition to the drug effect at the half maximum concentration (E_EC50_). **K-L**. The overall probability of cell death in G_1_ and S-G_2_ phase, respectively, for all five drug treatments. The arrow shows the shift from the control condition to the drug effect at the half maximum concentration (E_EC50_) for G_1_ and S-G_2_ phases.

To summarize the overall effect of each drug treatment, we compared the average phase durations and cell death probabilities inferred for each drug from the LCT model at the half maximum concentration (EC_50_) to the untreated control condition (**Fig. 2I-L**). The model inferred that lapatinib and palbociclib treatments lead to longer average G_1_ phase durations compared to untreated cells (**Fig. 2I-J**), a 10% higher chance of cell death in G_1_ phase for lapatinib-treated cells, and a slight chance of cell death in S-G_2_ after palbociclib treatment (**Fig. 2K-L**). The model also inferred that gemcitabine induces an increase in S-G_2_ durations and greater chance of cell death in S-G_2_ phase as compared to untreated cells (**Fig. 2G-H**). Finally, a 10% chance of cell death at the EC_50_ concentration (2.4 nM) was inferred in late G_2_ phase for cells treated with paclitaxel as compared to untreated controls (**Fig. 2J and Sup. Fig. 3J**).

### Analysis of single cell responses confirms model inferences and reveals drug-specific cell cycle phase effects

We developed model parameters from the average population response at each timepoint, which facilitates robust model development by leveraging information from a large number of cells. Importantly, as described above, the LCT model infers aspects of drug responses that can be quantified at the individual cell level—including cell cycle phase duration and cell cycle-specific death. We therefore tracked single cells in the image time course data to quantify cell cycle phase durations and also cell death events associated with specific drug treatments and concentrations (**Sup. Fig. 3B**). Quantification of cell death events also enables direct assessment of whether drug effects are cytotoxic or cytostatic. We analyzed the first complete cell cycle, which we reasoned would reveal early drug effects. We also quantified the relative fate outcomes for the progeny of cells at time 0H that underwent division, which provides insights into effects of drug treatment that are observed at later timepoints (**Sup. Fig. 3C**). As expected, in the untreated condition, most cells (93%) present at 0H underwent cell division. In contrast, at the highest lapatinib and gemcitabine doses, 32% and 61% of the cells present at time 0H failed to divide. Additionally, of the cells that did divide in these two conditions, only 10% underwent a second division. For both drugs, lower doses showed more modest changes in the fraction of cells that divided as compared to untreated. As described below, we compared these experimentally observed drug-induced cell cycle effects to those inferred by the LCT model.

The model inferred that the predominant lapatinib effect was to extend G_1_ durations from 22.3H in the untreated condition to 33.6H and 47.4H for 25 nM and 50 nM lapatinib, respectively (**Fig. 3A**). Experimentally, we observed that G_1_ durations increased after lapatinib (mean 26.2H and 32.5H with 25 nM and 50 nM lapatinib, respectively) (**Fig. 3A, B**). We also quantified an increase in the G_1_ duration variance showing that cells varied in their responsiveness to lapatinib (**Fig. 3B**). The model inferred minimal changes to S-G_2_ durations or cell death and we accordingly observed little change in S-G_2_ durations or cell death in the experimental data (**Fig. 3C,D**).

**Figure 3.**
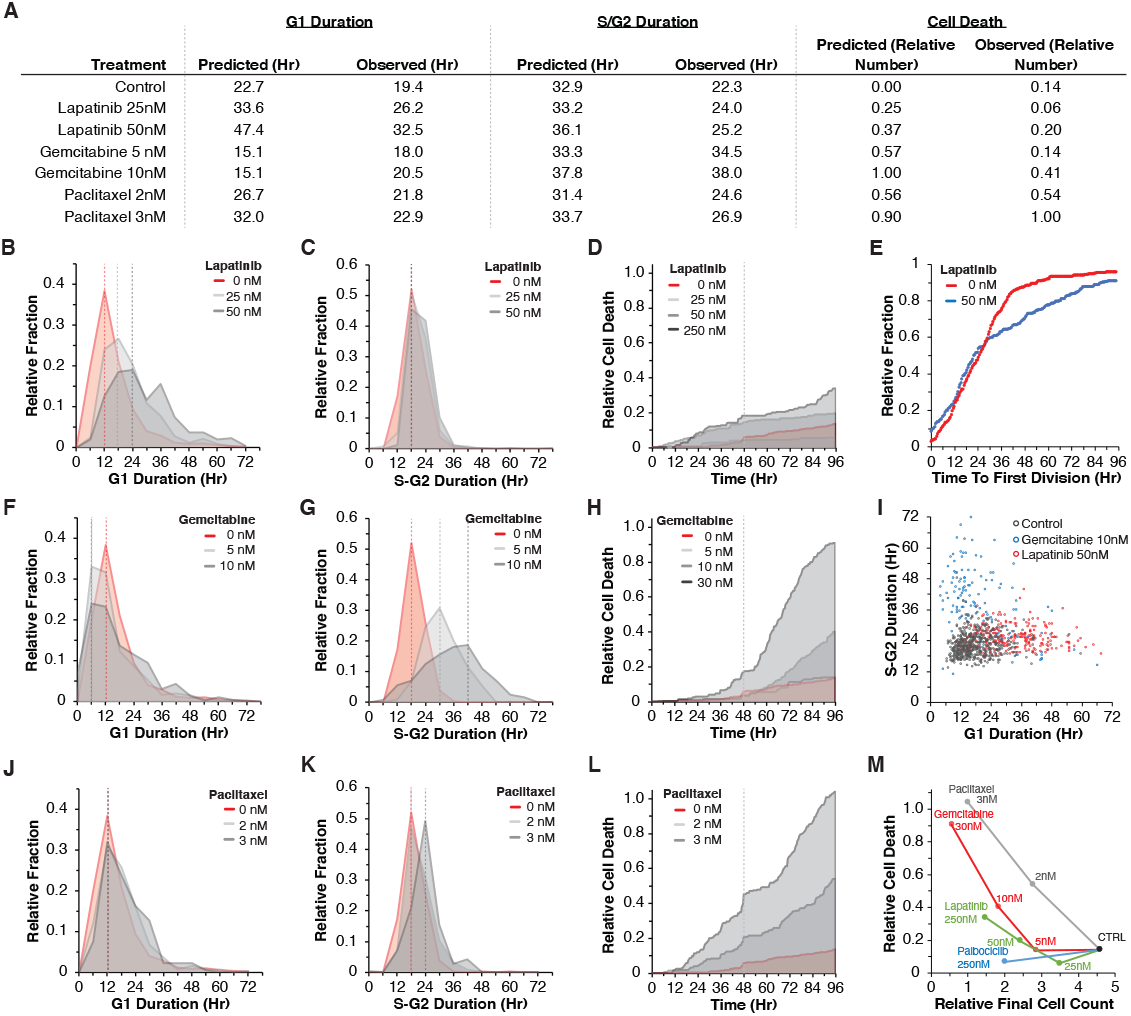
Analysis of single cell responses confirms model inferences and reveals drug-specific cell cycle phase effects. **A**. Quantification of cell cycle parameters as inferred by the model and observed experimentally (G_1_ and S-G_2_ durations and cell death). **B**. Distributions of G_1_ durations for cells that underwent one division in response to 0, 25, and 50 nM lapatinib. **C**. Distributions of S-G_2_ durations. **D**. Accumulated cell death across time. **E**. Time to first division for cells in the CTRL condition (red line) compared to 50 nM lapatinib (gray line). **F-H**. G_1_ and S-G_2_ distributions, and cell death accumulation in response to gemcitabine. **I**. G_1_ and S-G_2_ durations for the first complete cell cycle for all cells tracked in the control condition, in response to 100 nM lapatinib, and 10 nM gemcitabine. **J-L**. G_1_ and S-G_2_ distributions, and cell death accumulation in response to paclitaxel. **M**. Observed cell counts against cell deaths per drug.

The model inferred that oscillations in the percentage of G_1_ cells after lapatinib treatment arise from “waiting time “ effects in cell cycle progression (see **Fig. 2**). Waiting times, which can be modeled through distributions such as the gamma distribution, refer to the delay effect created by processes that are comprised of many sequential steps. To confirm the mechanism underlying this behavior at the single cell level, we examined various cell cycle measures and found a reduction in the fraction of cells undergoing their first division beginning around 24H (**Fig. 3E**). This observation, together with the lengthening of the subsequent G_1_ duration following cell division (**Fig. 3B**), can explain the cell cycle synchronization observed in the experimental data (see **Fig. 1**) and in the LCT model. At the start of the assay, cells in G_1_ are delayed in their time to division, while cells in S-G_2_ only become delayed at the onset of G_1_ following division. In effect, this creates two populations of cells with distinct timing in the induction of drug effects. We observed a similar effect after treatment with Palbociclib (**Sup. Fig. 3D**).

For gemcitabine, the model inferred a slight acceleration of G_1_ phases, which we also observed experimentally (**Fig. 3A,F**). The model inferred that S-G_2_ durations were extended following gemcitabine treatment, which we confirmed experimentally: S-G_2_ durations were extended from 22.3H to 34.5H with 5 nM and to 38H with 10 nM gemcitabine (**Fig. 3G**). Lastly, the model inferred an increase in the number of cell death events relative to the starting cell number, from 0 in control to 0.57 with 5 nM gemcitabine. At 10 nM gemcitabine, the model predicted 1.0 relative cell death events such that the number of cell death events across 96H was the same as the initial starting cell number (**Fig. 3A**). The experimentally observed values showed similar trends, though with more modest changes in cell numbers (0.14 and 0.41 relative cell numbers for 5 and 10 nM gemcitabine, respectively) (**Fig. 3H**). Overall, we observed similar trends in each of the parameters for gemcitabine treated cells as inferred by the model; modest differences were that the model inferred higher cell death and shorter extensions to S-G_2_ than we observed experimentally.

We also tested an assumption of the model that G_1_ and S-G_2_ phases are independent variables, which captures the idea that these cell cycle phases are independently regulated at the molecular level. We analyzed G_1_ versus S-G_2_ durations for individual cells in the control condition, 10 nM gemcitabine, and 50 nM lapatinib, and found a minimal correlation between G_1_ and S-G_2_ durations (**Fig. 3I**). These experimental observations confirm the implicit assumption of the model that G_1_ and S-G_2_ durations are uncorrelated.

Lastly, we evaluated model inferences for paclitaxel treatment. Consistent with our experimental observations, the model inferred minimal changes to G_1_ and S-G_2_ durations following treatment (**Fig. 3A,J,K**). At 2 nM paclitaxel, the model inferred 0.56 cell deaths relative to the starting cell numbers, and at 3 nM inferred 0.90 relative cell deaths (**Fig. 3A**, Methods). Experimentally, our observations were consistent with the values inferred by the model: we observed 0.54 and 1.00 relative cell deaths for 2 nM and 3 nM paclitaxel (**Fig. 3L**). To summarize the mechanisms that account for the observed changes in cell numbers due to paclitaxel treatment, we compared the number of cell death events against final cell counts for each of the other drugs. These data show the relative bias of paclitaxel toward inducing cell death, especially at 2 nM, compared to 5 nM gemcitabine and 50 nM lapatinib, which both resulted in similar final cell numbers (**Fig. 3M**).

Overall, the LCT model captured key observations about the cell cycle effects of each drug, which were confirmed by in-depth single-cell tracking of the experimental data.

### Drug-induced changes to cell cycle behavior generalize across a molecularly diverse panel of breast cancer cell lines

To assess the generalizability of our computational framework and experimental observations, we generated and tested three additional breast cancer cell lines from diverse molecular backgrounds^30^: 21MT1 (Basal subtype, HER2+), HCC1143 (Basal subtype, HER2-) and MDAMB157 (Claudin-low subtype, HER2-) (**Sup. Figs. 5-7**). Because these cell lines do not uniformly overexpress HER2, we additionally tested BEZ235 and trametinib, which respectively target PI3K and MEK, two growth factor pathways downstream from HER2. We observed dose-dependent reductions in cell numbers and also modulation of the percent of G_1_ cells following drug treatment. Importantly, similar to our findings for AU565 cells, we observed dynamic responses not captured by terminal endpoint readouts of cell viability (**Sup. Figs. 5-7, panels A-B**). We observed unique patterns, including: a delayed G_1_ enrichment from trametinib in 21MT1 cells (**Sup. Fig. 6**), a lack of G_1_ enrichment from palbociclib and BEZ235 in MDAMB157 cells (**Sup. Fig. 7**), and a dose-dependent bifurcation in G_1_ enrichment for doxorubicin in all three of the cell lines (**Sup. Figs. 5-7**).

Next, we tested our LCT model on each of the new cell lines. Comparison of model fits to experimental observations confirmed that our model could capture the dynamic responses observed across this panel of molecularly distinct cell lines, indicating the generalizability of our computational framework (**Sup. Figs. 5-7, panels C-E**). We analyzed the output of the LCT model, which inferred changes to cell cycle phase durations and cell death probabilities for drug-cell line pairs at the EC_50_ concentration (**Sup. Fig. 8**). The model inferred cell-line-specific changes to both G_1_ and G_2_ phases (**Sup. Fig. 8A,B**). For instance, 21MT1 were inferred to preferentially undergo G_1_ cell death after doxorubicin and paclitaxel treatments, at probabilities of 60% and 15%, respectively (**Sup. Fig. 8C**). The model inferred that HCC1143 cells arrest and die in S-G_2_ following paclitaxel or palbociclib treatment (**Sup. Fig. 8B,D**). MDA-MB-157 cells were inferred to become growth-arrested by drug treatment and to preferentially die in G_1_ phase (**Sup. Fig. 8C,D**). Overall, we confirmed that our computational framework was generalizable across several drugs and cell lines and could infer a range of drug treatment response behaviors.

### Responses to drug combinations are dependent on drug specific cell cycle and cell death effects

Durable and effective cancer treatments frequently require administration of multiple drugs; however identification of the principles underlying optimal drug combinations have been challenging^31^. Here, we tested the idea that our LCT model, which incorporates cell cycle effects, can be used to predict the impact of different drug combinations on cell cycle behavior and final cell numbers. We compared two strategies in accounting for drug combination effects. In the first, we combined drug effects on the rates of G_1_ and S-G_2_ progression using Bliss additivity and assumed the rates of cell death additively combined. In the second, we assumed an additive combination through use of the drug effects on overall cell numbers. To explore these predictions, we varied the dose of one drug in the two drug combination pair and analyzed responses to drug combinations that targeted either the same cell cycle phase (G_1_ and G_1_, or S-G_2_ and S-G_2_) or different cell cycle phases (G_1_ and S-G_2_).

First, we tested combining the rates for two G_1_ targeted drugs, such as lapatinib and palbociclib. The model predicted that effects on cell number would saturate around the initial starting cell number, indicating cytostatic effects of this drug combination (**Fig. 4A**). In contrast, drug combination effects based on cell numbers alone predicted a cytotoxic effect at higher drug concentrations, resulting in a reduction in cell numbers relative to the starting cell numbers. We tested these drug combinations experimentally and found a cytostatic effect at higher doses, which matches the model prediction based on combining rates of cell cycle progression (**Fig. 4A**).

**Figure 4.**
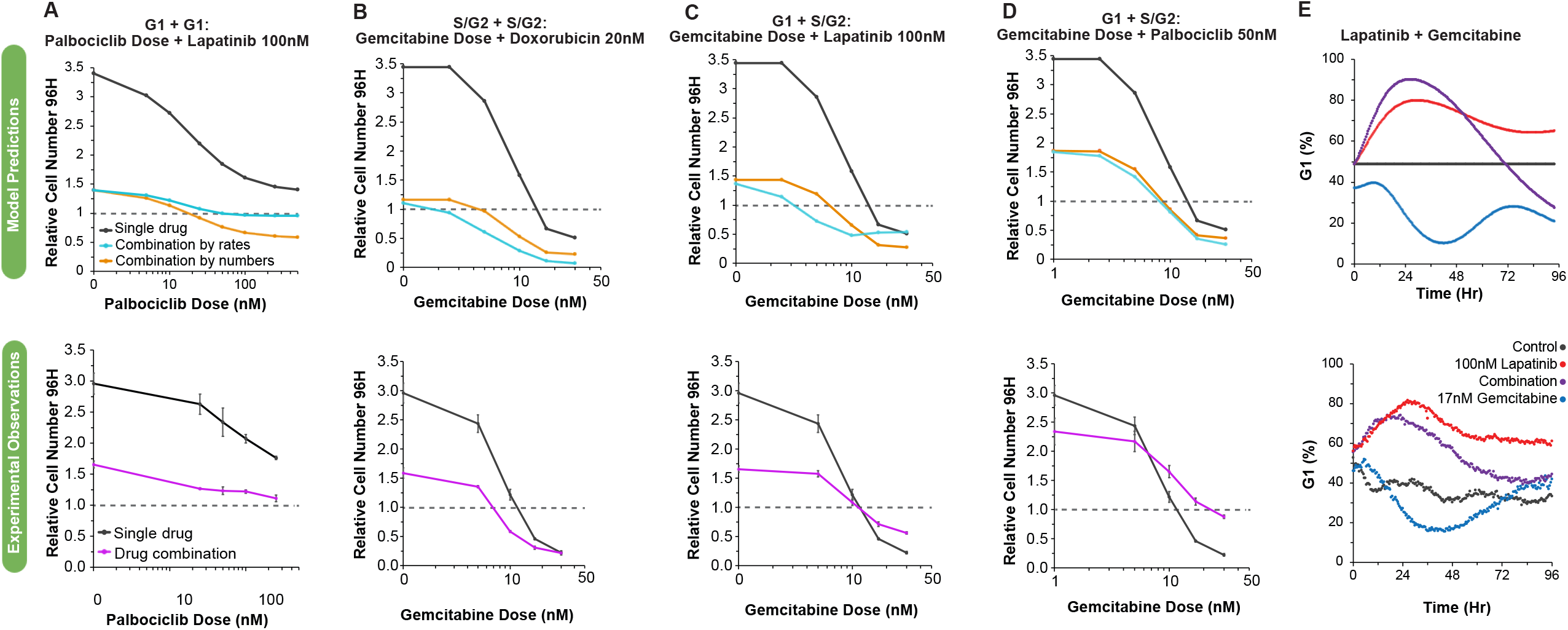
Responses to drug combinations are dependent on drug-specific effects on the cell cycle and cell death. **A-D**. Comparison between model predictions for single drug responses, or from drug combinations of Bliss additivity using cell numbers or model rates. **A**. Single drug responses for increasing doses of palbociclib and in combination with 100 nM lapatinib. **B-D**. Single drug responses for increasing doses of gemcitabine and in combination with 20 nM doxorubin, 100 nM lapatinib, 50 nM palbociclib. **E**. Model predictions for the percentage of cells in G_1_ phase for the control condition, 100 nM lapatinib, 17 nM gemcitabine, or the combination of lapatinib and gemcitabine. **E-H**. Comparison between the model predictions and experimental observations for the single drug responses and the drug combinations as described in panels A-D. Error bars show the standard error of the mean for three biological replicates.

Next, we analyzed predictions of gemcitabine combined with doxorubicin, which both extend S-G_2_ durations and induce cell death (see **Fig. 3A**). We found that combination predictions based on rates and cell counts both predicted a reduction in cell numbers relative to each drug on its own, which we also observed experimentally (**Fig. 4B**).

Lastly, we used the LCT model to examine the impact of combining two drugs that target different cell cycle phases, which mimics lapatinib (G_1_ effect) combined with gemcitabine (S phase effect). The cell cycle model predicted an antagonistic effect at higher doses, such that 30 nM gemcitabine combined with 100 nM lapatinib is expected to yield a similar final cell number as 30 nM gemcitabine on its own (**Fig. 4C**). Experimentally, we observed that three of the four lapatinib and gemcitabine combination doses showed an antagonistic impact on cell number as compared to gemcitabine alone indicating that combining these two drugs was actually counterproductive. These antagonistic effects of the combination held when lapatinib was replaced by palbociclib, which also impacted G_1_ durations (**Fig. 4D**). We examined the model predictions in more detail to gain insights into the underlying biological mechanisms driving these drug combination responses. The LCT model predicted that the G_1_ effect of lapatinib would initially dominate over the S-phase effects of gemcitabine, leading to an increased G_1_ proportion for the population, which was confirmed experimentally, thus mitigating the S-G2 effects of gemcitabine (**Fig. 4E**).

In summary, these data indicate that the cell cycle phase and cell death impacts of each drug in a pair are critical for determining the influence of single drugs on cell cycle behavior and that this information can be used for rational identification of drug combinations likely to be therapeutically beneficial.

## DISCUSSION

In this report, we link cell cycle regulatory mechanisms with drug-specific cell cycle effects to gain insights into cancer cell responses to individual drugs and drug combinations. To meld these ideas, we developed a combined experimental and modeling approach to measure cell dynamics and infer cell behavior. This combined approach revealed that assessment of temporal dynamics and cell behavior is critical to interpret and model drug-induced effects.

Importantly, assessment of the impacts of single agents on cell cycle behavior could be used to identify drug combinations likely to yield therapeutic benefits.

Recently, an in-depth analysis revealed that cell cycle phases in individual cells are uncorrelated and have durations that can be accurately modeled as an Erlang distribution (a special case of a gamma distribution)^32^. This observation indicates that the cell cycle can be viewed as a series of uncoupled, memoryless phases rather than a single process^10,33^. In this work, we found similar uncorrelated patterns in cell cycle phase responses after treatment with different anti-cancer drugs. This revealed multiple implications for assessing and modeling drug responses. First, viewing the cell cycle as a single process implies that cell behavior is immediately impacted upon drug treatment; however, we and others have reported that drug effects are often not observed until individual cells enter or approach a specific phase or checkpoint^34,35^. For instance, we found that cells were initially distributed across all phases of the cell cycle and that the addition of lapatinib, a G_1_-targeting drug, did not initially affect cells in S-G_2_ phase. This led to a partial cell cycle synchronization across the population and required the incorporation of a linear chain trick into our model to account for this dwell time. Additionally, the temporal dynamics of the therapeutic response were an important consideration for co-treatment with gemcitabine and lapatinib. If both drugs had immediate effects on cell behavior, we would expect that the G_1_ and S-G_2_ effects of each drug would counteract each other and lead to a constant ratio of cells in G_1_ phase. Instead, both experimentally and through model predictions, we found an initial G_1_ enrichment. This likely induced a secondary effect of reducing the relative time that each cell spent in S-phase, which further reduced gemcitabine sensitivity. This finding could explain the antagonistic effects on cell numbers that we and others have observed when combining gemcitabine with lapatinib or palbociclib^36,37^. We speculate that a synergistic effect on cell numbers could also arise by combining two drugs that target S-G_2_ phases, where each drug acts to extend the relative duration in which the other is effective. This general strategy could be used to identify optimal temporal scheduling of other drug combinations that induce different effects on the cell cycle.

A second implication of multiple independently regulated cell cycle processes relates to the concept of effect equivalence in drug combinations. This concept—that two drugs with independent targets can be used to identify drug synergy or drug antagonism—has predominantly focused on the cell number effect of each drug^2–5^. Our current work suggests that equivalence in effect may be better applied to rates of cell cycle phase progression and cell death. In our work, we found that lapatinib and palbociclib primarily impacted G_1_ phase with limited effects on cell death. In contrast, doxorubicin and gemcitabine extended S-G_2_ durations and induced cell death. These cell cycle and cell death effects were critical for gaining insights into the effect of drug combinations. For example, two cytostatic drugs, lapatinib and palbociclib, were additive up to doses that reached the maximum cytostatic effect, with further dose increases leading to only minor effects on cell numbers. In contrast, combining the two cytotoxic drugs led to increasingly cytotoxic responses across the full dose range. These results suggest that considering the cell cycle and cell death impacts of each drug is necessary to make predictions about the effects of their combinations and implies that this information could be used for the rational identification of effective drug combinations^38,39^.

Drug response measurements evaluated in the context of a mechanistic cell cycle model can reveal insights about the nature of drug response and resistance not immediately apparent from purely data-driven analyses. For instance, a model for the proliferation dynamics of cancer cells can separate the contribution of dividing, non-dividing, and dying cells^21^, revealing that the rates of cell death and entry into quiescence change with drug treatment. Previous computational models of cell cycle behavior have explored various ways in which cell cycle behavior might impact drug response but have struggled to identify experimental data amenable for model fitting and evaluation. For instance, others have appreciated that drugs do not affect the cell cycle uniformly and have therefore proposed computational models that partition the cell cycle into several independent steps, both with^10^ and without^33^ cell death effects. Modeling cell lifetimes as being hypo-exponentially distributed helps to explain the distribution of cell lifetimes within a population but does not connect these observations to known cell cycle stages^40^. In this report, we demonstrate that partitioning known cell cycle phases to account for their dwell time effects—and including experimentally observed drug effects like cell death—results in a modeling framework that can faithfully and mechanistically capture experimentally observed anti-cancer drug effects.

We applied our experimental approach and computational framework to examine dynamic drug-induced responses in a molecularly diverse set of breast cancer cell lines. In all cases, we observed that therapeutic inhibition induces a wide array of responses, indicating that the influence of therapies on cell cycle dynamics is a generalizable mechanism operable in a wide array of molecular backgrounds. Cancer cells treated with therapies may adopt new molecular programs associated with adaptive and acquired resistance, and indeed previous studies have demonstrated this principle in both model systems and patient samples^41^. We hypothesize that cells with acquired resistance may show distinct drug-induced cell cycle programs as compared to naïve cells and that our approach could be used to uncover the molecular mechanisms associated with adaptive resistance. Our study provides a blueprint for studying responses of diverse cell types—both normal and diseased—to a wide array of perturbations, including therapeutic inhibitors, growth factors, or genetic manipulation with CRISPRi/a. The resultant data could be used to adapt our computational framework to identify mechanisms of cell cycle control in different cellular contexts, microenvironmental conditions, or disease states.

While our model could explain many of the key observations in our experimental data, extensions of the model could further improve its generalizability and robustness. We partitioned the cell cycle into two observed phases, G_1_ and S-G_2_, which were further subdivided to explain the dwell time behavior of each phase. With improved reporter strategies^42^, we may be able to further subdivide these phases into constituent parts, which could help to localize the effect of a drug to a more specific portion of one cell cycle phase. Generalizations of the linear chain trick could be used to account for both subphases of varying passage rates, as well as heterogeneity in the rates of passage, such as would arise through cell-to-cell heterogeneity^29^. While the subdivisions within each cell cycle phase are phenomenological, it is tempting to imagine they represent mechanistic steps within each phase. Identifying how effects connect to actual biological events in the cell cycle would help identify opportunities for drug combinations. A practical challenge when using the model for drug combinations has been normalization between experiments. While cell number measurements are routinely normalized by dividing by a control, experiment-to-experiment variation in inferred rates requires additional consideration. A wider panel of experiments, across multiple cell lines, may help to tease apart variations associated with drugs, cell lines, or experiments. A final potential extension is considering the existence of phenotypically diverse subpopulations^43^. At the cost of additional complexity, one could employ several instances of the current model with transition probabilities between these states when the cells divide to simulate a heterogeneous population of cells.

## Summary

We observed that five commonly used cancer drugs each modulated cell numbers through distinct routes and with different temporal dynamics. By revealing how these drugs uniquely impacted cell fate, our model and analyses have implications for how different cancer drugs can be combined to maximize therapeutic impact. For instance, our results can identify drug combinations that modulate cell cycle effects in orthogonal ways or drug schedules that take advantage of the shift in cell cycle state of the overall population. In summary, these studies provide a map for understanding how cancer cells respond to treatment and how drugs may be combined and timed for maximal effect.

## ACKNOWLEDGEMENTS

These studies were supported by NIH research grants U54-CA209988, U54-HG008100, U01-CA215709, Prospect Creek Foundation, Jayne Koskinas Ted Giovanis Foundation for Health and Policy, and Breast Cancer Research Foundation. Cell sorting was performed by the OHSU Flow Cytometry Core, which is supported in part by the OHSU Knight Cancer Center Support Grant (P30CA069533) and an S10 instrument award (1S10OD028512).

## METHODS

### Creation of Stable Cell Lines

AU565 (ATCC CRL 2351) and MDAMB157 (ATCC HTB 24) cells were grown in DMEM supplemented with 10% FBS, HCC1143 (ATCC CRL 2321) cells were grown in RPMI supplemented with 10% FBS, and 21MT1 (generous gift from Kornelia Polyak) cells were grown in DMEM/F12 supplemented with 5% horse serum, 20 ng/ml rhEGF, 0.5 μg/ml hydrocortisone, 100 ng/ml cholera toxin, and 10 μg/ml insulin. The coding fragment for clover-HDHB was cloned in frame into a transposase expression plasmid modified to also express a nuclear localized mCherry^44^. The stable cell lines were created as previously described^45^ and selected for 7 days with 0.75 μg/ml puromycin. To mitigate a range of fluorescent signals from transfection, HCC1143 and 21MT1 cells were sorted at OHSU ‘s Flow Cytometry Core and cells with a medium intensity clover-HDHB signal and a high intensity NLS-mCherry signal were selected for drug dose response experiments. In all cases, cells were validated by STR profiling (LabCorp) and tested negative for mycoplasma.

### Drug Dose Response Protocol

AU565 cells were plated at a density of 25,000 cells per well into 24-well Falcon plates (Corning #353047). 24H after plating the media was exchanged with Fluorobrite media supplemented with 10% FBS, glutamine, and penicillin-streptomycin. Cells were then treated with dose-escalation: lapatinib (Selleckchem #S1028), gemcitabine (#S1149), paclitaxel (#S1150), doxorubicin (#S1208), palbociclibb (#S1116), BEZ235 (#S1009), and trametinib (#S2673). After drug addition, plates were imaged every 30 minutes for 96H using phase, GFP, and RFP imaging channels with an IncuCyte S3. For single drug treatments of AU565 cells only, at 48H the media was replaced in all wells including the control wells, and fresh media and drug were added. Four equally-spaced image locations per well and three biological replicates were collected.

MDAMB157, HCC1143, and 21MT1 cell lines were transferred to and maintained in a base of either Fluorobrite media and 1x GlutaMAX or mixed Fluorobrite/F12 media and 0.5x GlutaMAX along with their corresponding supplements for no less than one week before performing the drug dose response protocol. MDAMB157 and HCC1143 cells were plated at a density of 25,000 cells per well, while the larger 21MT1 cells were plated at a density of 5,000 cells per well into 24-well Falcon plates (Corning #353047). 24H after plating the media was exchanged with fresh Fluorobrite media as indicated per cell line. Cells were then treated with dose-escalation: BEZ235, gemcitabine, paclitaxel, doxorubicin, palbociclib, and trametinib. After drug addition, plates were imaged every 2 hours for 96H using phase, GFP, and RFP imaging channels with the IncuCyte S3. Four equally-spaced image locations per well and three biological replicates were collected.

### Image Analysis

To analyze AU565 image data, phase, GFP, and RFP images were overlaid and collated into single files using FIJI^46^, then segmented into three classes (nuclei, background, debris) using a manually trained classifier in Ilastik^47^. The segmented nuclear masks from Ilastik and the IncuCyte GFP images were used to count the number of nuclei in each image with Cell Profiler^48^. Additionally, using the same images (nuclear masks from Ilastik and GFP cell cycle reporter images) cell cycle phase was determined by taking the mean fluorescence in the nucleus compared to the mean fluorescence in a 5-pixel ring surrounding the nucleus, excluding background pixels. A threshold was then manually set for the ratio of nuclear fluorescence to cytoplasmic fluorescence and cells with values below the threshold were defined as being in G1 and cells with values above the threshold were defined as being in S/G2 phase^48^.

To manually track AU565 cells and identify drug-induced changes operable in single cells, GFP image sequences were registered using the FIJI plug-in ‘StackReg ‘. Individual cells present in the first image and their progeny were followed to identify the time of G_1_ transition, cell death, and cell division using the plug-in mTrackJ^49^. We excluded cells that were binucleated, had abnormally large nuclei, or were near the image border where complete lineages could not be tracked. The G_1_ transition was defined as the last frame before the nuclear intensity of the cell cycle reporter was below the level of the cytoplasm. Assessment of cell death enabled disentangling of cytostatic and cytotoxic drug effects.

We used the following approach for automated analysis of HCC1143, 21MT1 and MDAMB157 cell lines. Image registration was performed on the red channel nuclear marker image stack using the python skimage phase_cross_correlation function to correct translations. Image stacks were cropped to their common areas and individual cells were segmented with the Cellpose LC2 model trained on phase and nuclear images from the untreated and highest drug concentration treatments^50^. Nuclei were segmented with the Cellpose cyto2 model on the nuclear channel. To associate nuclei across the image stack, to identify progeny after mitosis, and to identify cell death events we used Loeffler tracking^51^ with the default parameters of delta_t = 3 and roi_size = 2. We created cytoplasm masks by subtracting the nuclear masks from the cell masks and applied these masks to the green channel cell cycle reporter images using the python skimage function regionprops_table. To assign cells to G1 or S/G2 states, we computed the ratios between the cytoplasm and nuclear cell cycle reporter. k-means clustering of the ratios observed in cells in the untreated condition was used to establish a per-plate threshold between cell cycle states.

The quantitated cell-level data was mean summarized to the population level for each image and to assess cell counts and G1 cell cycle state proportion. The cell counts were normalized to the mean of the counts of the first three images. The cell count dose response curves were normalized to the control by dividing each drug cell count by the control cell count at the same time slice.

### Core Model

To identify the dynamics of the AU565 cancer cell population in response to compounds, we built a system of ordinary differential equations (ODEs) with two states: G_1_, and S-G_2_. Cells transition from G_1_ to S-G_2_ phase, and then vice versa when doubling. Cell death can occur in either phase with phase-specific death rates. S and G_2_ phases are combined as our reporter cannot distinguish them. From single-cell tracking, we identified that G_1_ and S-G_2_ phase time-intervals are gamma-distributed. Based on this observation, we employed the linear chain trick (LCT)^28^ to capture these waiting time distributions. We broke down each phase into a series of sequential sub-phases and derived the system of mean-field ordinary differentials. Each sub-phase is represented as a single state variable within the differential equation system. The total number of cells in each phase is the sum of the cell numbers in each sub-phase. Furthermore, to account for the non-uniform effect of the drugs over each cell cycle phase, we divided G_1_ and S-G_2_ into 4 parts each, such that the effect of a drug can be distinguishable at the beginning, middle, or the end of the phases.

The mean-field system of ODEs is:

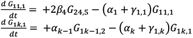

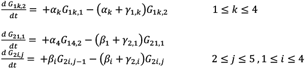

The parameters of the model include progression rates through G_1_ phase, *α*, and S-G_2_ phase, *β*, and death rates in each of the G_1_ phase, *γ*_1_, and S-G_2_ phase, *γ*_2_. Cells at the end of the S-G_2_ phase divide and give birth to two cells at G_1_ phase. Because each phase is divided into 4 parts, each part of G_1_ contains 2 sub-phases, and each part of S-G_2_ contains 5 sub-phases.

The model was implemented in Julia v1.5.3. The differential equations were solved by the matrix exponential. As the data was measured with equal spacing, we pre-calculated the transition matrix between timesteps.

### Dose Response Relationship

We assumed that the progression and death rates in G_1_ and S-G_2_ that form the quantified drug effects on the population follow a Hill function:

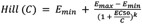

where the *EC*_50_indicates the half-maximal drug effect concentration, *E*_*min*_ the value of the rate parameter in the absence of drug, *E*_*max*_ the rate parameter at infinite concentration, and *k* the steepness of the dose-response curve. Given these parameters and the drug concentration (*C*) we then calculated the specific rate parameters for that treatment.

### Exponential Model

To show the benefit of our LCT model, we employed a commonly used exponential model to fit to the G_1_ and S-G_2_ cell numbers and showed that the exponential model cannot capture the dynamics of the data. The parameters were the same as the mean-field model.

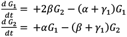

### Model Fitting

The data included the percentage of cells in G_1_ phase and the total number of cells normalized to the cell numbers at the initial time point. We assumed 1 starting cell at 0H and calculated the number of cells in G_1_ and S-G_2_ phase over time. The Savitzky-Golay filter was used to smooth the data. Three replicates of each experiment were averaged, and the average was used for the purpose of fitting.

The number of G_1_ and S-G_2_ subphases, and the parameters in the absence of drug were shared across all drugs and concentrations. The sum of squared error was used as the cost function value and was calculated between the cell numbers predicted by the model and the average cell numbers of three replicates, over all time points, concentrations, and drugs tested. This cost function was then minimized using the default adaptive differential evolution optimizer from the BlackBoxOptim.jl Julia package, version 0.5.0.

To characterize the identifiability of our fit parameters we conducted a local sensitivity analysis. To do so, we calculated the cost function while varying each parameter from 0.1 to 10 times the optimal value, holding all the other parameters at their optimum (results not shown). We observed that all parameters were identifiably constrained by this analysis.

### Calculating relative number of cell deaths and average phase durations

We evaluated the number of dead cells at 96H relative to the starting cell number at 0H. This formed the observed relative cell death numbers reported in Figure 3A. To calculate the corresponding cell death values inferred from the model, we calculated the predicted number of cells at each phase part (G_11_, G_12_, G_13_, G_14_, G_21_, G_22_, G_23_, G_24_) separately, and multiplied them by their individual death rates at all time points. This provides the number of dead cells at each phase part at each time point. The sum of cell numbers died in each phase part provides the total cell death counts at each time point, *n*(*t*. Figure 2C-D show the accumulated dead cells across time for lapatinib and gemcitabine treatments which was calculated by summing over the cell death counts, *n*(*t*), across time from 0 to each timepoint, *T*. Calculating for 96H results in the total cell death normalized by the initial cell numbers, 1, this value refers to the relative predicted cell death number reported in Figure 3A.

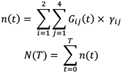

The average phase durations 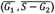 from the model were calculated using the progression rates. The G_1_ phase has 8 subphases which is divided into 4 parts that results in 2 phases per part. S-G_2_ phase has 20 subphases divided into 4 parts that results in 5 subphases in each part. Each phase part has a unique parameter for cell death and phase progression rate. The average phase duration will be given by the following expressions, derived by recognizing that the time in each part is gamma-distributed with a shape parameter equal to the number of subphases.

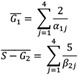

### Predicting Drug Combinations

Bliss independence was used to calculate the predicted effect of drug combinations. Assuming *E*_*a*_and *E*_*b*_ to be the saturable, quantified effects of drugs *a* and *b*, the expected combined effect would be:

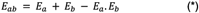

For death effects, we added the effects of each drug to find the death effect of the drug combination:

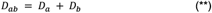

The Bliss relationship requires that data first be scaled to be between 0 and 1, and then scaled back after the interaction calculation:

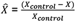

This measure is usually used as a baseline to decide whether the combination of two drugs is synergistic or antagonistic. Here we used Bliss in two ways: (1) on the progression rate parameters to simulate the model predictions of drug combinations; and (2) on cell numbers to serve as a baseline approach to calculate drug combination effects, as is commonly used. In the first case, we use Bliss additivity on the cell cycle progression rates (*) to find the set of progression parameters representing the combined treatment and assume that the death effects are only additive because the cell death process is not saturable (**). The combination parameters for all the eight concentrations for all pairs of drugs were calculated and then converted back to their original units. Next, we simulated the cell numbers using these parameters. In the baseline case, we used the cell numbers in the control condition to normalize the cell number measurements and then converted the cell numbers back to their original scale. This was used as a benchmark reference.

## Data Availability

Data was deposited to the Image Data Resource (https://idr.openmicroscopy.org) under accession number idr0119. All other data and analyses used in this study are available from the corresponding author upon reasonable request.

## Code Availability

The code and analysis can be found at https://github.com/meyer-lab/DrugResponseModel.jl. Code for the automated segmentation and tracking is at https://github.com/markdane/CellTracking.

## FIGURE LEGENDS

**Figure S1. Individual replicates for AU565 drug responses show similar temporal dynamics and drug-induced changes to cell cycle**. Panels show relative cell numbers and S-G_2_ normalized cell numbers for lapatinib (**A**), gemcitabine (**B**), paclitaxel (**C**), palbociclib (**D**), and doxorubicin (**E**) treatments for three biological replicates. Five drug concentrations (gray lines) and untreated control (red line) are plotted.

**Figure S2. An exponential cell cycle model without incorporating delay times fails to capture the dynamics of drug response. A**. The transition diagram for a simple dynamical model with 2 phases (G_1_ and S-G_2_) and without the LCT. *α* and *β*, are the transition rates from G_1_ to S-G_2_ and vice versa, *γ*_1_ and *γ*_2_ are the death rates in G_1_ and S-G_2_, respectively. **(B-E)**.

Exponential cell cycle model simulations of G_1_ and S-G_2_ cell numbers over time for control and 5 concentrations of lapatinib **(B-C)** and gemcitabine **(D-E)** (solid lines), respectively, overlayed with the average of three experimental replicates (dashed lines).

**Figure S3. Analysis of single cell tracking data reveals drug-specific cell cycle phase effects in AU565 cells. A**. Lineage trees of 25 lineages across 96H for various drug treatments. Tracks are colored coded based on cell cycle phase: gray indicates G_1_ and red indicates S-G_2_ phase. Track splitting indicates mitosis, and track ending prior to 96H corresponds to apoptosis. **B**. Quantification of cell outcomes (division, apoptosis, still present at end of experiment) for cells from the first and second generations treated with lapatinib, gemcitabine, or paclitaxel. **C**. Gamma distribution of G_1_ and S-G_2_ phase durations for cells in control condition with sample size of 520 and 514 for G_1_ and S-G_2_ phases, respectively. **D**. Lineage trees for 25 lineages across 96H after treatment with Palbociclib.

**Figure S4. A dynamical model of the cell cycle captures the dynamics of drug response. A-F**. G_1_ and S-G_2_ cell numbers overtime, respectively, for the control and treatment at 5 concentrations (solid lines) for doxorubicin (**A-B**) paclitaxel (**C-D**), and palbociclib (**E-F**) overlayed with the average of three corresponding experimental replicates (dashed lines). **G-L**. The average phase durations in G_1_ and S-G_2_ phases for selected drug treatments. The arrow shows the shift from the control condition to the drug effect at the half maximum concentration (E_EC50_).

**Figure S6. The introduced dynamical model captures the cell cycle dynamics of drug response in 21MT1 cell line. A,B**. Experimentally observed drug-induced changes to cell numbers (**A**) and G_1_ cell cycle phase proportion (**B**) after dose-escalation treatment with a panel of inhibitors. **C,D**. G_1_ and S-G_2_ fits overtime, respectively, for the untreated and treatment at 5 concentrations (solid lines) overlayed with the average of three corresponding experimental replicates (dashed lines) for 6 drug treatments. **E**. Inferred accumulated dead cells over time for 6 drug treatments.

**Figure S5. The introduced dynamical model captures the cell cycle dynamics of drug response in TNBC cell line HCC1143. A,B**. Experimentally observed drug-induced changes to cell numbers (**A**) and G_1_ cell cycle phase proportion (**B**) after dose-escalation treatment with a panel of inhibitors. **C,D**. G_1_ and S-G_2_ fits overtime, respectively, for the untreated and treatment at 5 concentrations (solid lines) overlayed with the average of three corresponding experimental replicates (dashed lines) for 6 drug treatments. **E**. Inferred accumulated dead cells over time for 6 drug treatments.

**Figure S7. The introduced dynamical model captures the cell cycle dynamics of drug response in TNBC cell line MDA-MB-175. A,B**. Experimentally observed drug-induced changes to cell numbers (**A**) and G_1_ cell cycle phase proportion (**B**) after dose-escalation treatment with a panel of inhibitors. **C,D**. G_1_ and S-G_2_ fits overtime, respectively, for the untreated and treatment at 5 concentrations (solid lines) overlayed with the average of three corresponding experimental replicates (dashed lines) for 6 drug treatments. **E**. Inferred accumulated dead cells over time for 6 drug treatments.

**Figure S8. Summary of inferred cell cycle drug effects at half maximum concentration compared to untreated. A-B**. The average phase durations in G_1_ **(A)** and S-G_2_ **(B)** phases for HCC1143 (blue), 21MT1 (olive) and MDA-MB-157 (pink) treated with paclitaxel, palbociclib, trametinib, BEZ235, doxorubicin, and gemcitabine. The dashed lines show the average phase duration at untreated for each cell line. **C-D**. The cell death probability in G_1_ **(C)** and S-G_2_ **(D)** phases for HCC1143 (blue), 21MT1 (olive) and MDA-MB-157 (pink) treated with the same panel of drugs. The arrows show the quantity of increase or decrease in the effects from untreated to the half maximal concentration (E_EC50_).

