## Supplemental Table 1 for "Analysis and modeling of cancer drug responses using cell cycle phase-specific rate effects"

| Symbol | Units | Explanation |
| --- | --- | --- |
| $\alpha_{1}$ | 1/cell/hr | Cells progression rate through the first quarter of G1. |
| $\alpha_{2}$ | 1/cell/hr | Cells progression rate through the second quarter of G1. |
| $\alpha_{3}$ | 1/cell/hr | Cells progression rate through the third quarter of G1. |
| $\alpha_{4}$ | 1/cell/hr | Cells progression rate through the fourth quarter of G1. |
| $\beta_{1}$ | 1/cell/hr | Cells progression rate through the first quarter of S/G2. |
| $\beta_{2}$ | 1/cell/hr | Cells progression rate through the second quarter of S/G2. |
| $\beta_{3}$ | 1/cell/hr | Cells progression rate through the third quarter of S/G2. |
| $\beta_{4}$ | 1/cell/hr | Cells progression rate through the fourth quarter of S/G2. |
| $\gamma_{11}$ | 1/cell/hr | Cells rate of death within the first quarter of G1. |
| $\gamma_{12}$ | 1/cell/hr | Cells rate of death within the second quarter of G1. |
| $\gamma_{13}$ | 1/cell/hr | Cells rate of death within the third quarter of G1. |
| $\gamma_{14}$ | 1/cell/hr | Cells rate of death within the fourth quarter of G1. |
| $\gamma_{21}$ | 1/cell/hr | Cells rate of death within the first quarter of S/G2. |
| $\gamma_{22}$ | 1/cell/hr | Cells rate of death within the second quarter of S/G2. |
| $\gamma_{23}$ | 1/cell/hr | Cells rate of death within the third quarter of S/G2. |
| $\gamma_{24}$ | 1/cell/hr | Cells rate of death within the fourth quarter of S/G2. |

Supplementary Table 1. Explanation of the rate parameters within the ODE model.
