## Supplemental Table 2 for "Analysis and modeling of cancer drug responses using cell cycle phase-specific rate effects"

| Cell line | Drug | EC50 concentration [nM] |
| --- | --- | --- |
| HCC1143 | Paclitaxel | 1.46 |
| HCC1143 | Palbociclib | 86.46 |
| HCC1143 | Trametinib | 3.71 |
| HCC1143 | BEZ235 | 34.94 |
| HCC1143 | Doxorubicin | 159.16 |
| HCC1143 | Gemcitabine | 0.74 |
| 21MT1 | Paclitaxel | 1.55 |
| 21MT1 | Palbociclib | 19.1 |
| 21MT1 | Trametinib | 220.8 |
| 21MT1 | BEZ235 | 12.52 |
| 21MT1 | Doxorubicin | 51.68 |
| 21MT1 | Gemcitabine | 1.51 |
| MDAMB157 | Paclitaxel | 2.01 |
| MDAMB157 | Palbociclib | 3594.01 |
| MDAMB157 | Trametinib | 658.20 |
| MDAMB157 | BEZ235 | 10.90 |
| MDAMB157 | Doxorubicin | 247.58 |
| MDAMB157 | Gemcitabine | 0.64 |

Supplementary Table 2. The concentration at half maximal effect for each cell line and treatment.
