## Supplementary figures and images for "Analysis and modeling of cancer drug responses using cell cycle phase-specific rate effects"

### Supplementary Figure 1

#### Supp. Figure 1

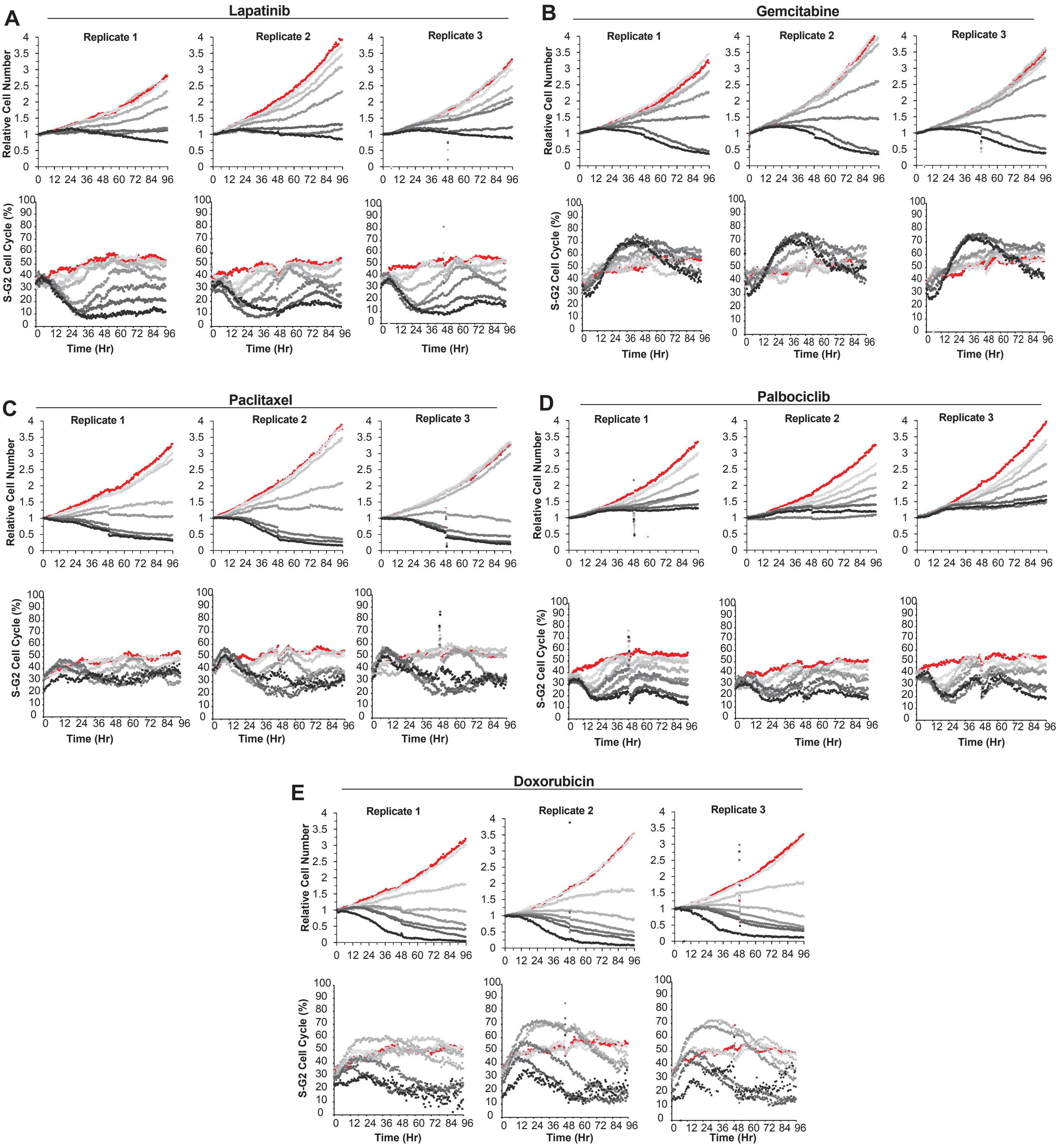

### Supplementary Figure 2

#### Supp. Figure 2

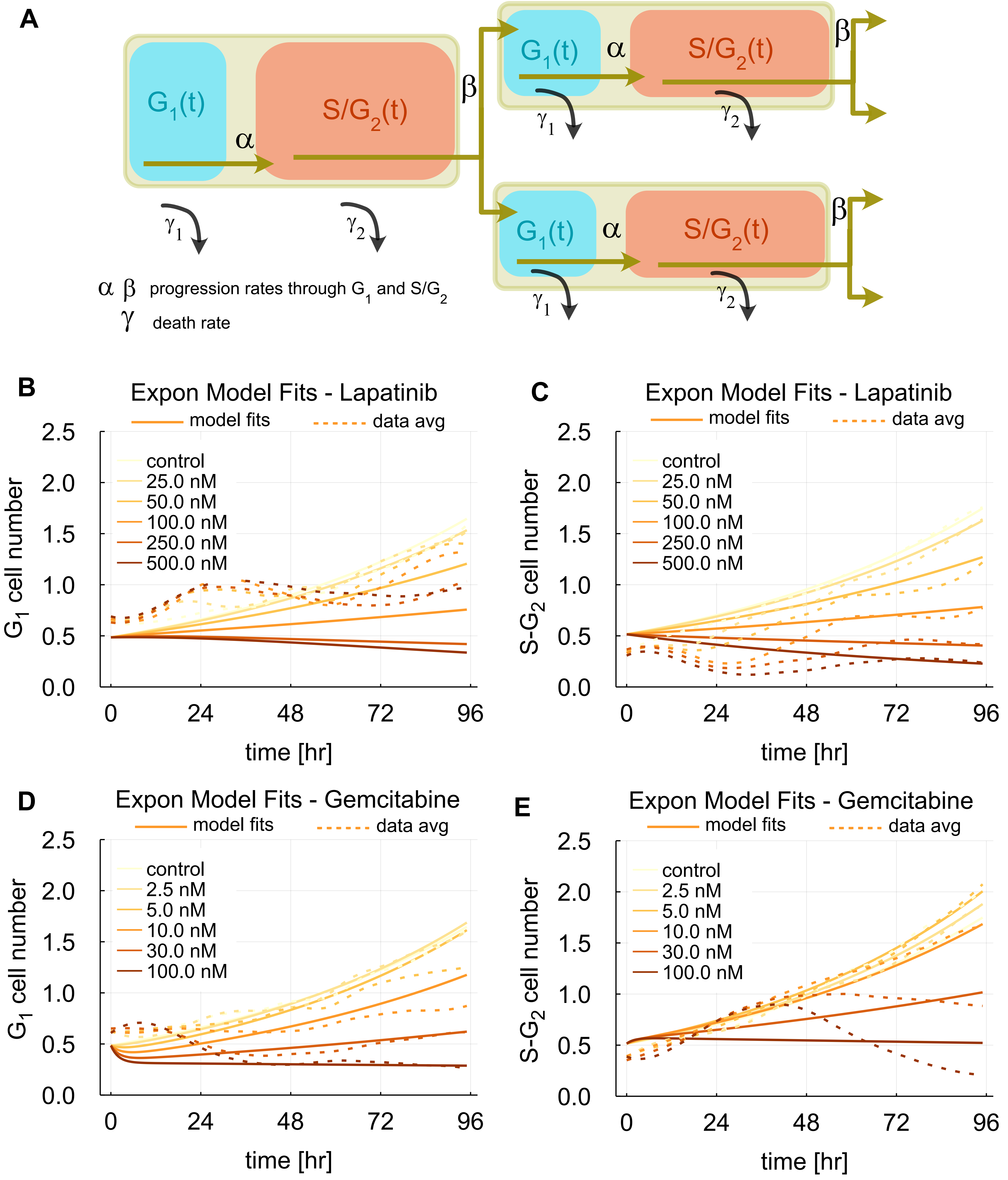

### Supplementary Figure 3

#### Supp Figure 3

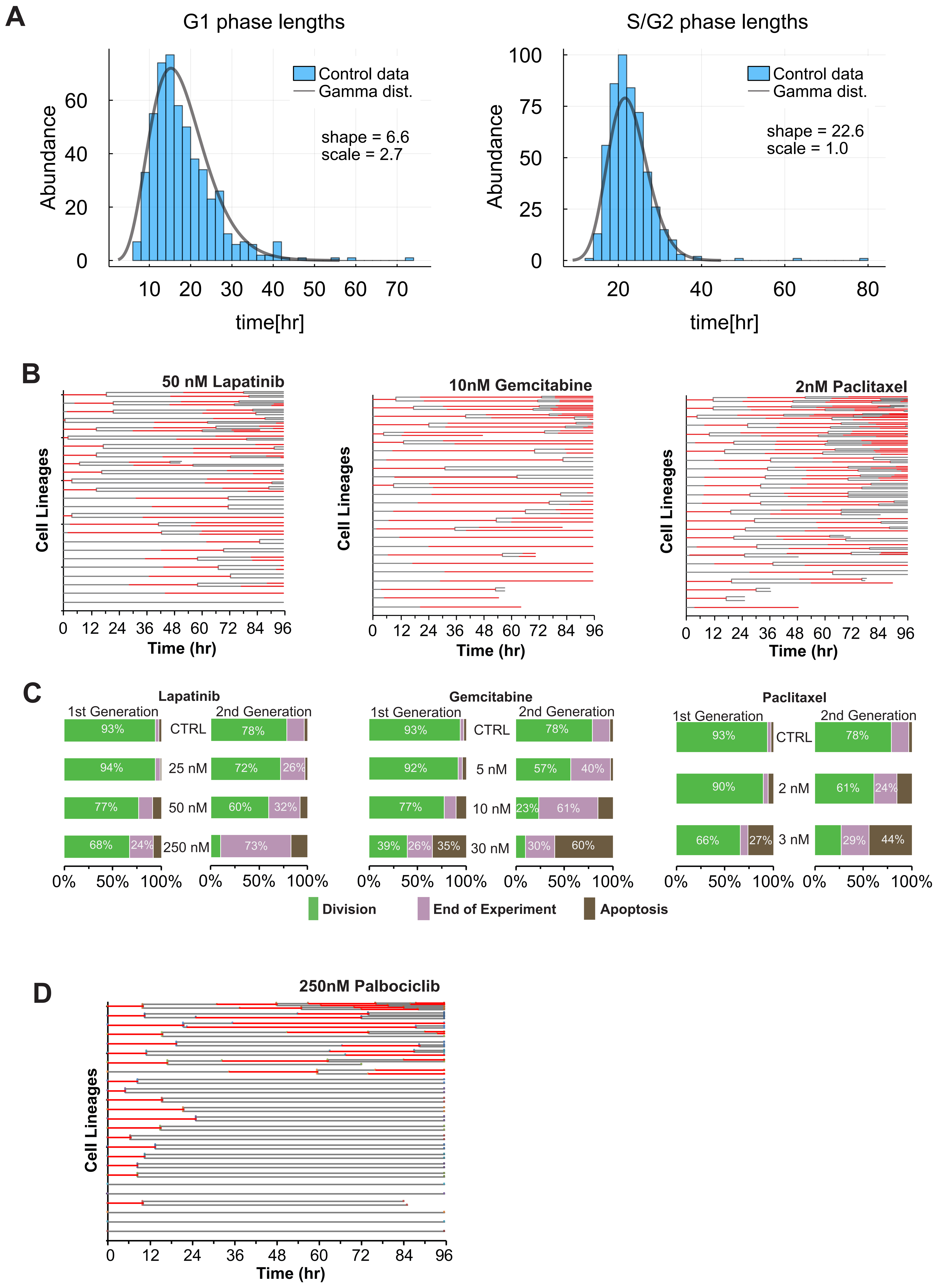

### Supplementary Figure 4

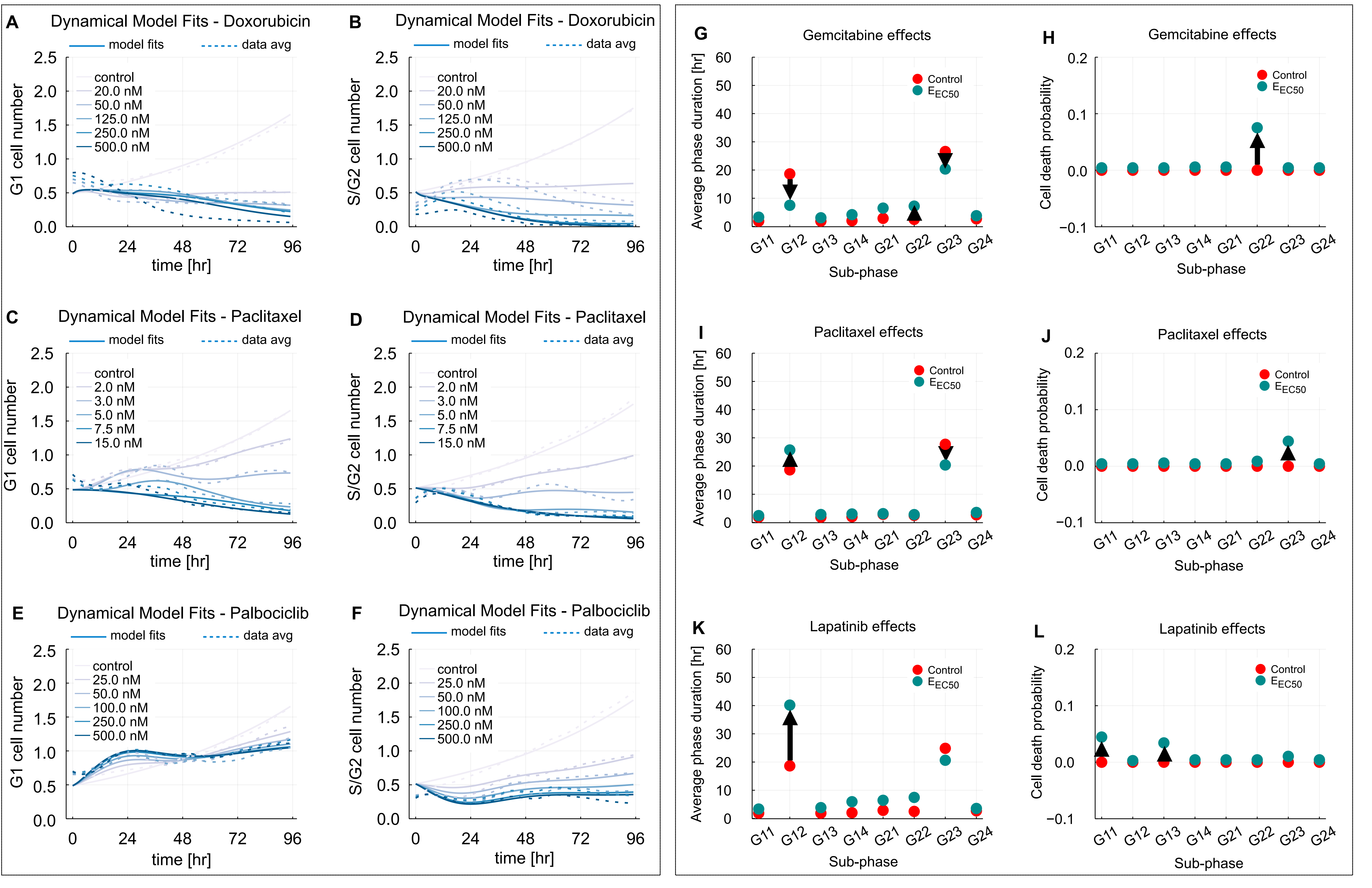

### Supplementary Figure 5

#### Supp. Figure 5

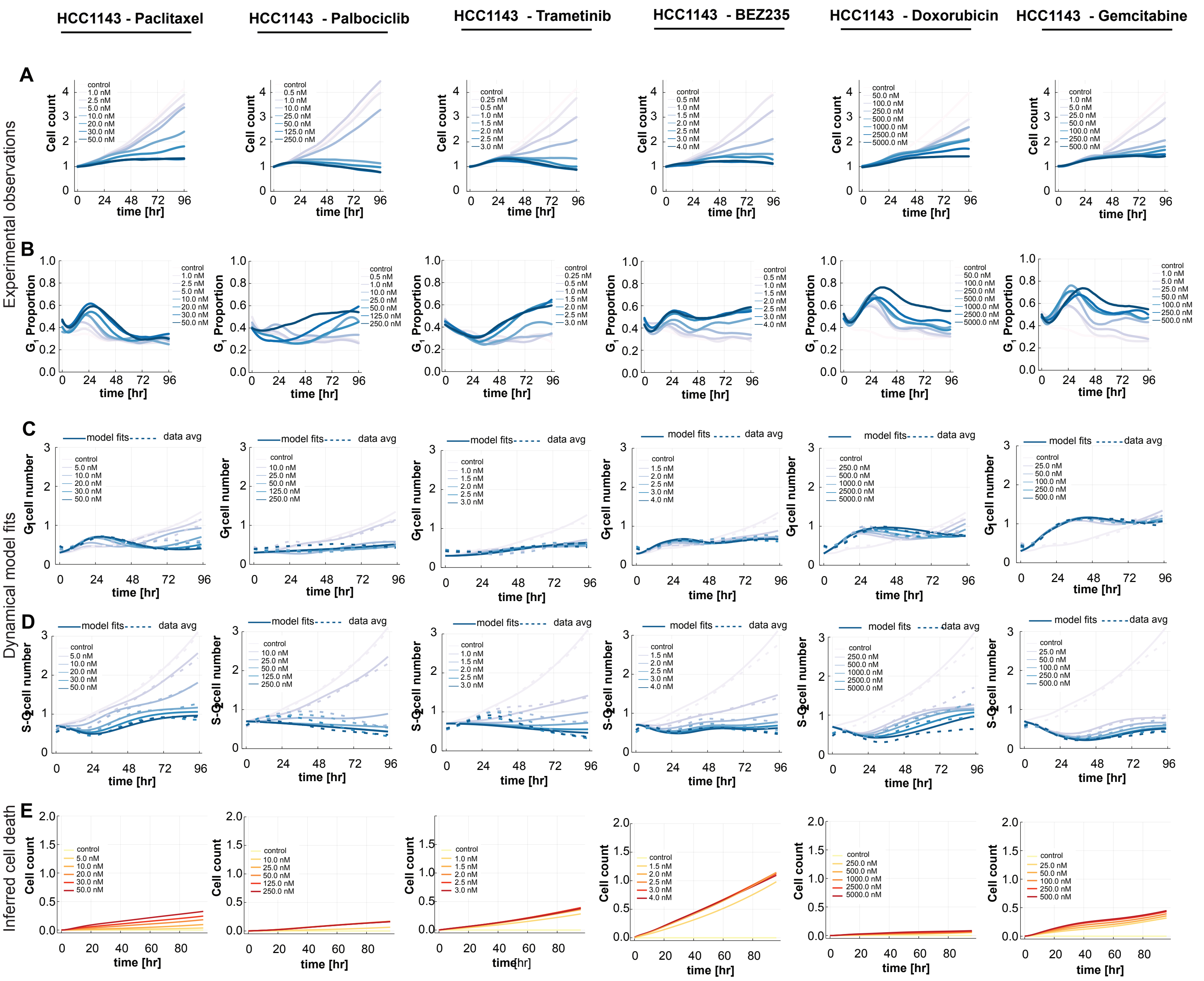

### Supplementary Figure 6

#### Supp. Figure 6

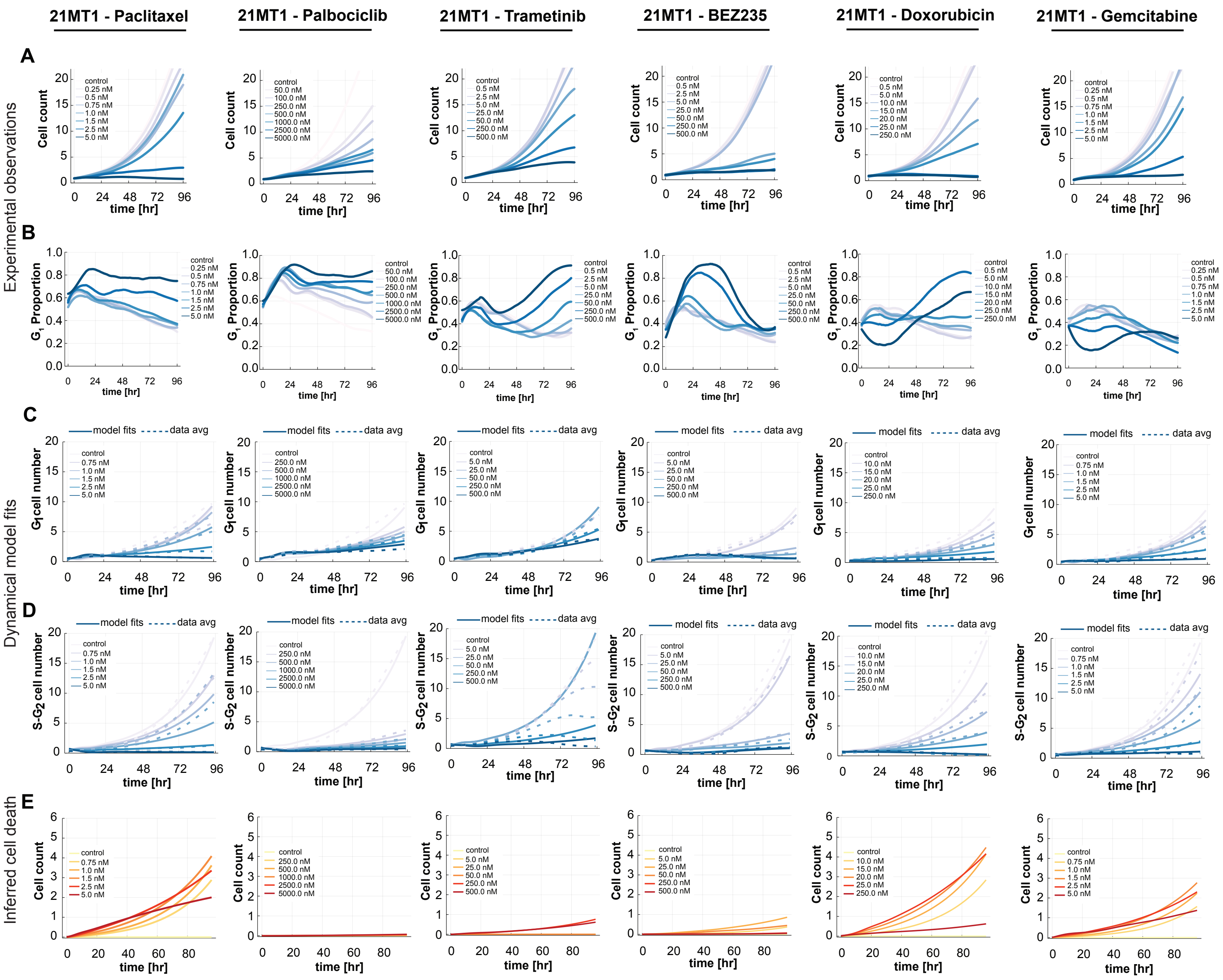

### Supplementary Figure 7

#### Supp. Figure 7

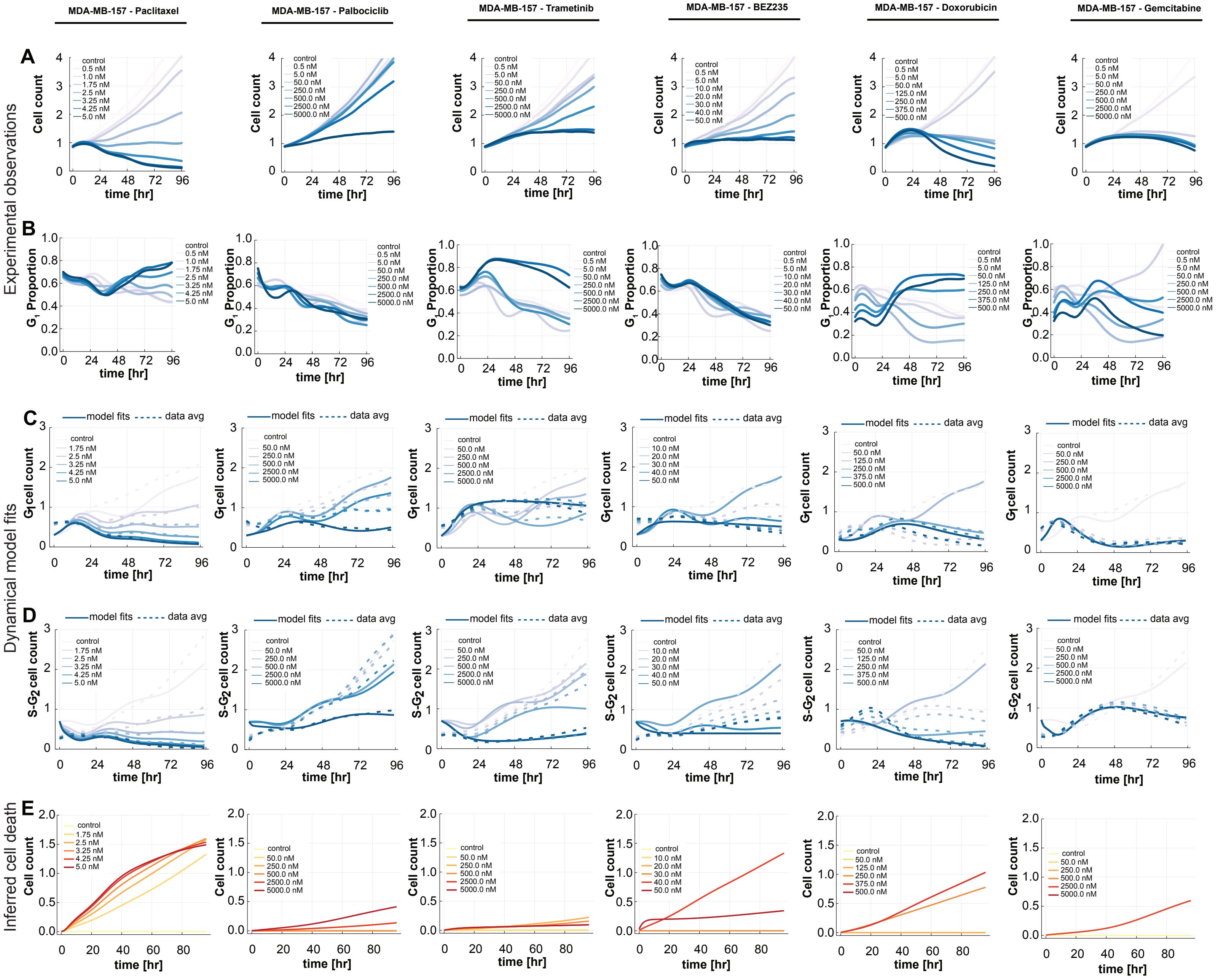
